# Extended period of selection for antimicrobial resistance due to persistency of antimicrobials in broilers

**DOI:** 10.1101/2024.04.18.590069

**Authors:** Aram F. Swinkels, Bjorn J.A. Berendsen, Egil A.J. Fischer, Aldert L. Zomer, Jaap A. Wagenaar

## Abstract

Antimicrobials can select for antimicrobial resistant bacteria. After treatment the active compound is excreted through urine and faeces. As some antimicrobials are chemically stable and very persistent, recirculation of sub-inhibitory concentrations of antimicrobials may occur due to coprophagic behaviour of animals such as chickens.

The persistence of three antimicrobials over time and their potential effects on antimicrobial resistance were determined in four groups of broilers. Groups were left untreated (control) or were treated with amoxicillin (non-persistent), doxycycline or enrofloxacin (persistent). Antimicrobials were extracted from the faecal samples and concentrations were measured by LC-MS/MS. We determined the resistome genotypically using shotgun metagenomics and phenotypically by using *Escherichia coli* as indicator microorganism.

Up to 37 days after treatment, persistent antimicrobials (doxycycline and enrofloxacin) had concentrations in faeces equal to or higher than the minimal selective concentration (MSC), in contrast to the non-persistent (amoxicillin) treatment. The amoxicillin treatment showed a significant difference (p ≤ 0.01 and p ≤ 0.0001) in the genotypic resistance only directly after treatment. On the other hand, the doxycycline treatment showed approximately 52% increase in phenotypic and a significant difference (p ≤ 0.05 and p ≤ 0.0001) in genotypic resistance throughout the trial. Furthermore, the enrofloxacin treatment resulted in a complete enrofloxacin-resistant *E. coli* population but the quantity of resistance genes was similar to the control group, likely because resistance is mediated by point mutations.

Based on our findings, we suggest that persistency of antimicrobials should be taken into consideration in the assessment of priority classification of antimicrobials in livestock.

## Introduction

Antimicrobial compounds are crucial and lifesaving medicines which can be applied to treat infections with a bacterial pathogen and preventively for surgeries or organ transplantations. ^1–3^ Unfortunately growing bacterial resistance against antimicrobial compounds has become an increasing problem during the last decades. ^4^ The rising development of antimicrobial resistance (AMR) has been considered as one of the most serious health threats for humans and animals by organizations as the World Health Organization (WHO), the Food and Agriculture Organization of the UN (FAO) and the World Organisation for Animal Health (WOAH). ^3,5,6^ The excessive use of antimicrobials in livestock is contributing to the emergence of resistant bacteria. ^7^ Measures to reduce antimicrobial usage in livestock are important to reduce the exposure to antimicrobial concentrations that select for resistant bacteria.

In terms of reducing the worldwide antimicrobial usage, antimicrobial stewardship programmes have been established. ^8,9^ These programmes consist of guidelines to reduce the use of antimicrobials or to stimulate prudent usage of antimicrobials in livestock. ^10–12^ For example the European Medicine Agency (EMA) has classified antimicrobial compounds into four different categories; A (avoid), B (restrict), C (caution) and D (prudence). ^13,14^ This categorization of antimicrobial compounds has been established to reduce potential consequences for public health as an result of increased AMR. For that reason antimicrobials in the categorization A are not licensed for animal species. The category B include antimicrobial compounds which have been listed as medically important antimicrobials (MIAs) for the human health sector by the WHO. ^15,16^ Furthermore, the category C should only be considered when there are no clinically effective alternatives in category D. ^15^

Based on the classification of the EMA, antimicrobials are selected and applied by veterinarians to limit the selection of AMR to MIAs. However, in this categorization, only direct selection during treatment is considered. Studies have shown that there is an extensive difference in half-life properties between antimicrobial compounds ranging from hours to over a year. This implies that, for specific antimicrobials, selection for resistant bacteria can be maintained long after the application and withdrawal time. ^17,18^ Especially in poultry the persistency of antimicrobial compounds is of concern due to coprophagic behaviour, which induces recirculation of the antimicrobial compounds thereby re-exposing the gut microbiome to residual concentrations that may be above the minimal selective concentration (MSC) for a longer period which results in a prolonged selective pressure. ^19–22^ Since persistence is not included in the assessment of the classification of antimicrobials it could be a missed opportunity for reducing AMR. To elaborate, to our knowledge, no studies have been conducted which show a relation between persistence of antimicrobial residues in combination with resistance levels.

In our study we measured the temporal antimicrobial residue concentration in faecal droppings and caecal material after treatment of broilers. Parallel to this we investigated the prevalence of AMR in *Escherichia coli* isolates and the resistance genes in resistome data after cessation of the antimicrobial treatment. An animal trial was conducted in which groups of broilers were treated with amoxicillin (non-persistent), doxycycline and enrofloxacin (both persistent). Antimicrobials were extracted from the faecal samples and analysed by liquid chromatography coupled to tandem mass spectrometry (LC-MS/MS) and were compared to the minimum selective concentrations (MSC) determined by Swinkels et al. (manuscript under revision). We performed phenotypic resistance assays by plating *E. coli* isolates, determining the proportion of wildtype vs non-wildtype. Additionally, we also quantified the read depth of resistance genes present in the resistome to investigate resistance genes besides the indicator organism *E. coli*. Lastly, we studied whether the antimicrobial residuals had any effect on the composition of the microbiome.

## Material and methods

### Housing of the broilers, experimental setup and sample collection

A total of 158 broilers (commercial Ross 308) were included in this trail and were transported from the hatchery to the animal facility (Utrecht University, Utrecht, The Netherlands) on the day of hatching (day 0 of their life). The broilers were weighed, tagged and housed in one pen (6 m^2^) to equilibrate their microbiome. After four days, caeca were collected after euthanizing 12 broilers. Subsequently, the remaining broilers were randomly divided into different treatments groups (day 0 of the experiment) and treatment started. The groups were treated with amoxicillin (non-persistent), doxycycline or enrofloxacin (both persistent) and an untreated control group. Each group was divided into three subgroups containing 12 broilers. The groups stayed in separate stables with hygiene rules to prevent carry-over of antimicrobials and bacteria. The antimicrobial treatment lasted four days with concentrations used in practice according to the SPC. ^23–25^ Afterwards faecal droppings were collected individually from the broilers placed in carton boxes with paper at the bottom. The faecal droppings were homogenized to obtain pooled samples and a fraction was stored at −80°C for antimicrobial analysis. This was repeated at day 6, 12, 19, and 26 after the start of the treatment. On day 37, after treatment, broilers were euthanized, and caecal material was collected. The samples were transported to the lab and further processed.

### Ethics of experimentation

Broilers were observed daily for the presence of clinical signs, abnormal behaviour and mortality. The study protocol was approved by the Dutch Central Authority for Scientific Procedures on Animals and the Animal Experiments Committee of Utrecht University (Utrecht, the Netherlands) under registration number AVD10800202114909 and all procedures were done in full compliance with all legislation. A power calculation was conducted to estimate the total number of broilers needed for the experiment (Supplementary data, Zenodo repository).

The size of the pens of the subgroups was set to 2 m^2^ according to the legal regulations in appendix III of 2010/63/EU ^26^ to not exceed the kilograms per square meter. Loss of broilers in the first week of the experiment due to health issues was expected, resulting in 12 broilers per subgroup. Solely female broilers were selected to reduce weight increase. ^27^

### Phenotypic resistance

From every subgroup and time point a swab was used to inoculate the pooled samples on three MacConkey plates. Next, 24 single colonies were picked by a pipette tip after overnight culture at 37°C. The colonies were transferred to a single well in a 96-wells plate with 100 µl LB medium per well. The 72 colonies per treatment and time point in one 96-wells plate were transferred to squared MacConkey plates with ECOFF (epidemiological cut-off values) concentrations of amoxicillin (8 mg/L), doxycycline (4 mg/L), enrofloxacin (0.125 mg/L) and a control plate without antimicrobials via a stamp. Next, non-wildtype colonies were scored after overnight incubation at 37 °C. An *E. coli* colony was scored as non-wildtype (further referred to as resistant) if it was able to grow on the selection plate with the respective antimicrobial. We calculated *E. coli* resistance by comparing growth on control and selection plates.

### Shotgun metagenomics

DNA was extracted according to the EFFORT protocol, DNA concentrations were measured with a Qubit. ^28,29^ Illumina sequencing was performed using Illumina NovaSeq 6000 (Useq, Utrecht sequencing facility) with a maximum read length of 2 x 150 bp. Libraries were prepared with Illumina Nextera XT DNA Library Preparation Kit according to the manufacturers protocol. ^30^ Afterwards, the reads were trimmed with trim galore (v0.6.4_dev) and the quality was assessed with FastQC (v0.11.4). The reads were analysed for the taxonomic classification by Kraken2 and the abundance of the DNA sequences was computed with Bracken. ^31,32^ The output was summarized into a biomfile using kraken2biom and was analysed with the R programme packages Phyloseq (v1.36.0), Microviz (v0.9.1) and Microbiome (v1.14.0) by which the species composition, alpha and beta diversity were estimated from a rarefied phyloseq object. ^33,34^ Finally the resistome was investigated by using KMA V 1.4.2 utilizing the Resfinder database with a minimum of 80% gene coverage. ^35,36^ The reads were first normalised for gene length and displayed as sequence depth per gigabase of sequencing data. Sequence data was uploaded in the SRA under accession PRJEB73721.

### Antimicrobial analysis of the faecal samples

The faeces samples were stored at −80 °C until extraction of the antimicrobial compounds. The samples were analysed according to the procedure described by Berendsen et. al 2018. ^18^ In short, from the samples, approximately 1 gram was weighed into a 50 ml propylene (PP) centrifuge tube and internal standard solution was added. The antimicrobial compounds were extracted by 4 ml of McIlvain-EDTA buffer and 1 ml acetonitrile. Afterwards 2 ml of lead acetate solution was added to remove excessive proteins. After centrifugation, the extract was transferred to a clean test tube and diluted by 13 ml 0.2 M EDTA. A Phenomenex (Torrance, CA, USA) Strata-X RP 200 mg/6 ml reversed phase solid phase extraction (SPE) cartridge was conditioned with 5 ml MeOH and 5 ml water. Then the complete extract was applied onto the SPE cartridge and washed with 5 ml of water and dried by applying vacuum for 5 min. The antimicrobial compounds were eluted by adding 5 ml methanol to the cartridges. The eluate was then evaporated at 40 °C under nitrogen. The residue was redissolved in 200 µl methanol and 300 µl water was added before transferring the extract into an LC-MS/MS sample vial.

The LC system consisted of a Shimadzu UFLC XR (Milford, MA, USA) model Acquity with a Phenomenex Kinetex C18 analytical column of 1.7 µ C18 100A, 100*2.1 mm, placed in a column oven at 40 °C. The gradient profile with a flow rate of 0.3 ml/min is shown in Table 1. The injection volume was 5 μl. Detection was carried out by LC–MS/MS using a Sciex (Framingham, MA, USA) Q-Trap 6500 mass spectrometer in the positive electrospray ionization (ESI) mode. The antimicrobials were fragmented using collision induced dissociation (N2) and the scheduled Selected Reaction Monitoring (SRM) transitions (20 s window) as described in Berendsen et al. (2015). ^37^ Data was processed using Multiquant software V2.1.1 (Sciex).

**Table 1.**
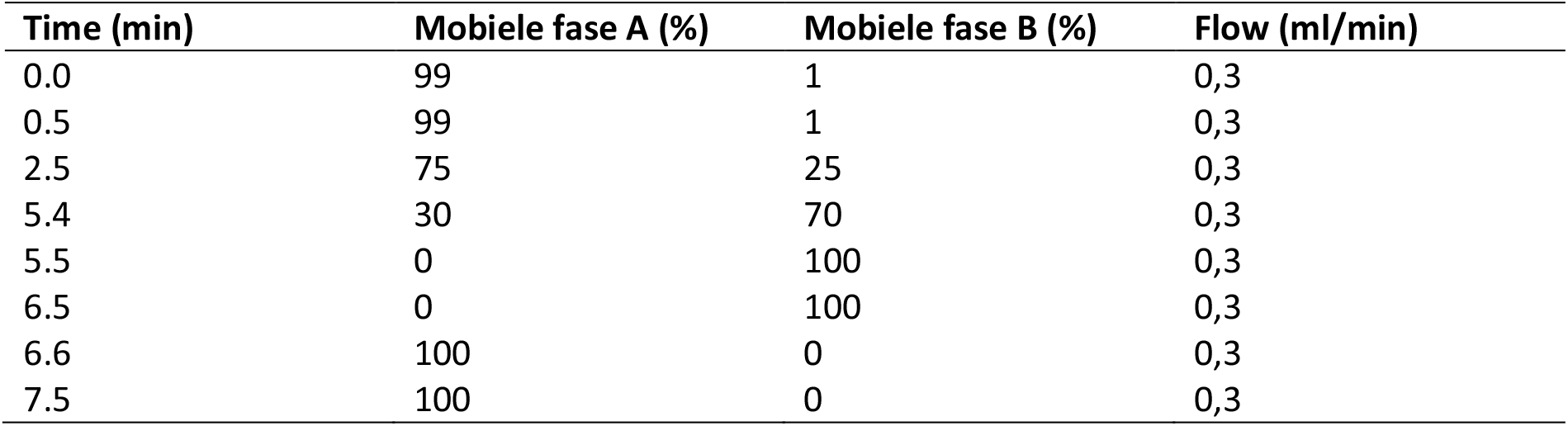
Gradient profile used in the LC-MS/MS method.

### Statistical analysis

We used R (version 4.1.0) and the package lme4 (version 1.1-29) for statistical calculations. We used logistic mixed effects and linear mixed effects models. Full models included Treatment and Time and the interaction between Treatment and Time as fixed effects and Pen as random effect. A logistic regression model was fitted with an interaction term for time after treatment and treatment. This was necessary due to inflated standard errors caused by counts of 0 (no resistance) or 24 (all resistance). Models without Time and /or Treatment were compared to the full model. The best model was selected based on the AIC. For the resistome data, we performed a post hoc Tukey test to determine whether the treatment groups at the different time points were significantly different from the control group. The microbiome diversity was measured with Shannon index for the alpha diversity and Bray-Curtis distance for the beta diversity. All the R-script and data files can be found in the supplementary data, Zenodo repository 10.5281/zenodo.10720991.

## Results

### Antimicrobial residues extracted from the faecal samples before, during and after treatment

The concentrations of the antimicrobial residues extracted from the faecal samples are shown in Figure 1. At time point 0 and in the control group we did not measure any antimicrobial compounds. For the non-persistent antimicrobial amoxicillin we did not find concentrations which reached the MSC. Only at six days after the start of the treatment we measured amoxicillin at 0.05 mg/kg, however this was below the MSC. On the contrary, the persistent antimicrobials doxycycline and enrofloxacin showed relatively high concentrations after treatment. For doxycycline the concentrations were above or within the range of the MSC up till 26 days. Enrofloxacin concentrations were far above the MSC range after treatment and was still in the MSC range at the time of slaughter. These results suggest that doxycycline and enrofloxacin remain selectively effective beyond application.

**Figure 1.**
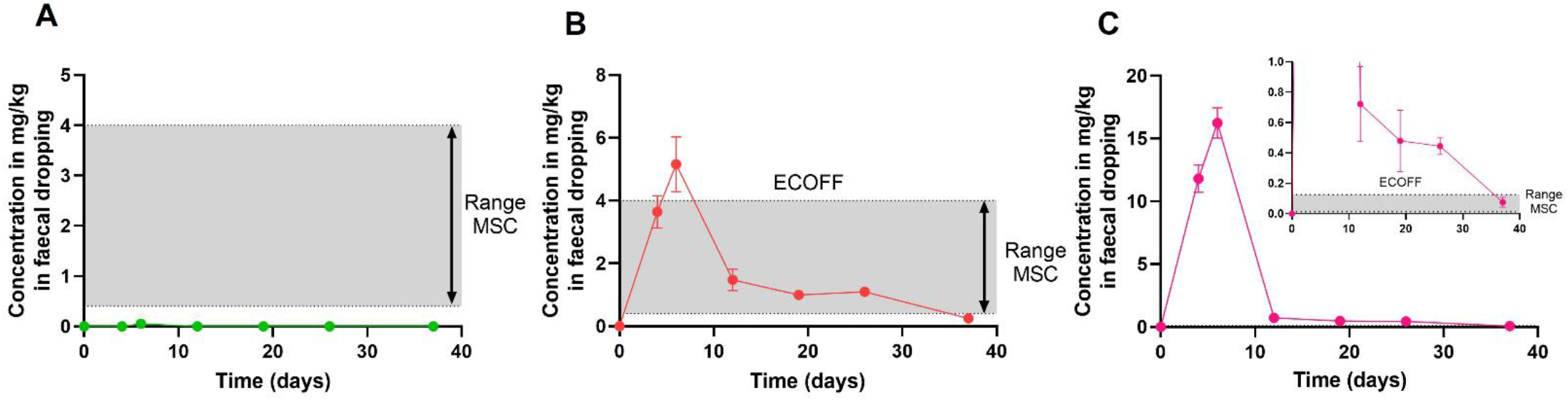
Antimicrobial extraction in faecal droppings of broilers treated with different antimicrobials. **A** amoxicillin, **B** doxycycline and **C** enrofloxacin. In the graphs the minimal selective concentration (MSC) are also displayed, determined by Swinkels et al. (manuscript under revision). The MSC was determined by concentrations with a 10 fold difference, therefore the established MSC is displayed as a range between the lowest and highest possible concentration in the figures. The epidemiological cut-off values (ECOFF) are distinguishing bacteria from wildtype and non-wildtype (in this study called resistant). The datapoints are presented by the mean and standard error of the mean.

### Quantifying phenotypic resistance of *E. coli* isolates over time

The *E. coli* isolates which are isolated over time in the different treatment groups are shown in Figure 2. We observed that the *E. coli* isolates from the amoxicillin treatment started with high proportion of resistant *E. coli* followed by a declining trend comparable to the control group. For the doxycycline treatment we observed a steep increase directly after treatment and a slight decline at time point 12 days, however the majority of the *E. coli* isolates remained resistant to doxycycline up until the slaughter age while in the control group the number of resistant *E. coli* remained limited. For the enrofloxacin treatment we found almost exclusively enrofloxacin resistant *E. coli* isolates which is in strong contrast to the control group where we observed only a few *E. coli* isolates that harboured resistance towards enrofloxacin. The data were best explained (lowest AIC) with a model containing both time after treatment and treatment.

**Figure 2.**
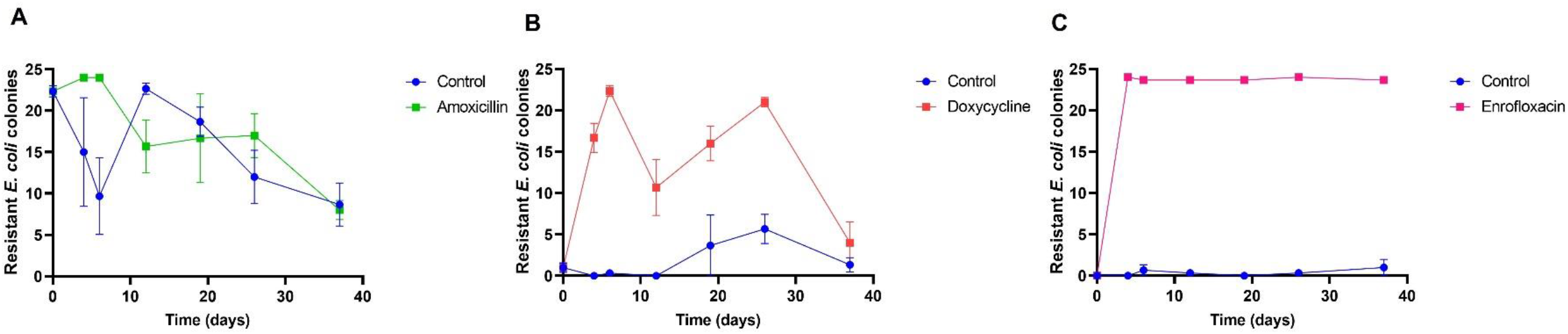
Phenotypic resistance of E. coli colonies. The graph are showing the resistance of E. coli towards the antimicrobial which a group of broilers is treated with compared to the control group. Graph A amoxicillin treatment compared with the control group, graph B doxycycline compared with the control group and graph C enrofloxacin compared with the control group. The datapoints are presented by the mean and standard error of the mean.

### Quantifying resistance genes in faecal droppings

We determined the resistome to quantify resistance genes present in the different treatment groups (Figure 3).

**Figure 3.**
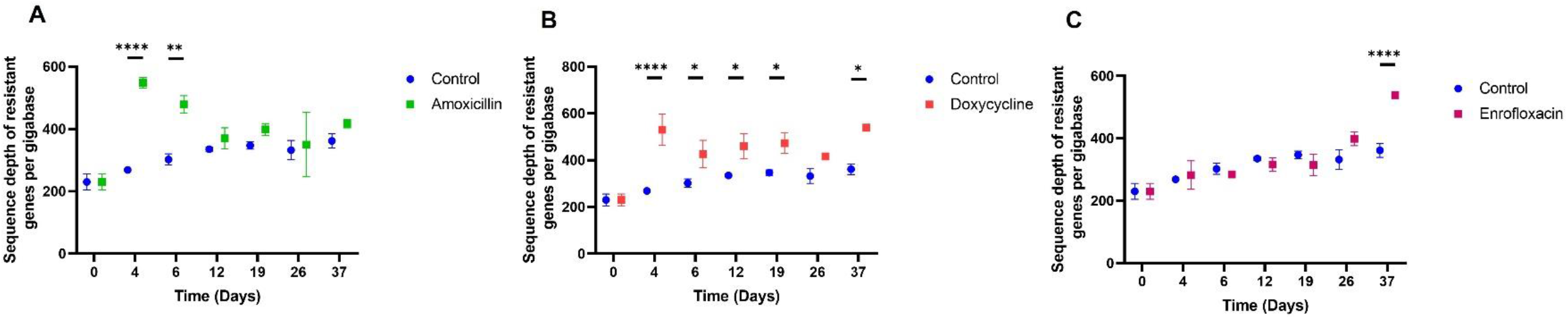
Number of resistance genes in the resistome in sequence depth per gigabase over time. In graph A the amoxicillin treatment is shown compared to the control group, in graph B the doxycycline treatment with the control group and in graph C the enrofloxacin treatment with the control group. The data points are presented by the mean and standard error of the mean. * p ≤ 0.05, ** p ≤ 0.01, **** p ≤ 0.0001.

Regarding amoxicillin treatment, we observed more resistant genes immediately after the start of treatment (day 4) and at day 6 than in the control group (amoxicillin – control day 4 p < 0.0001, day 6 p = 0.0023). However by day 12 of the treatment, the quantity of resistant genes in the treatment group approached levels similar to those in the control group. Doxycycline treatment led to a similar trend, with increased resistance genes at day 4 and day 6 compared to the control group (doxycycline – control day 4 p < 0.0001, day 6 p = 0.0161). However, the difference in resistant genes between the doxycycline-treated group and the control group persisted even at day 12 (p = 0.0156) and day 19 (p = 0.0149). Notably, the first decline in resistant genes within the doxycycline group, approaching control group levels, was observed 26 days after treatment initiation. As for enrofloxacin treatment, its impact on resistant genes was closer to that of the control group, especially up to day 19. On day 26, a minor increase in resistant genes was observed but no significant difference. It was only after slaughter in the caecal that a significant difference was observed between the enrofloxacin-treated group and the control group (enrofloxacin – control day 37 p < 0.0001).

### Microbial composition determination after treatment with antimicrobials

We studied the composition of the microbiome to determine if exposure to the residual antimicrobials after treatment had an effect on the microbial composition (Figure 4). The most noticeable change was the *E. coli* abundance in the microbiota of the treated groups. The enrofloxacin treatment resulted in a strong reduction of *E. coli* to almost 0% at day 4, afterwards it returned slowly in the microbiome. Furthermore, the same was observed for the doxycycline treatment as it reduced *E. coli* to 10% at day 4 and returned to levels comparable to the control group. Contrarily, the amoxicillin treated group favoured *E. coli* at day 4 and its abundance declined over time. Notably, at day 12, the control group had almost 90% *E. coli* abundance.

**Figure 4.**
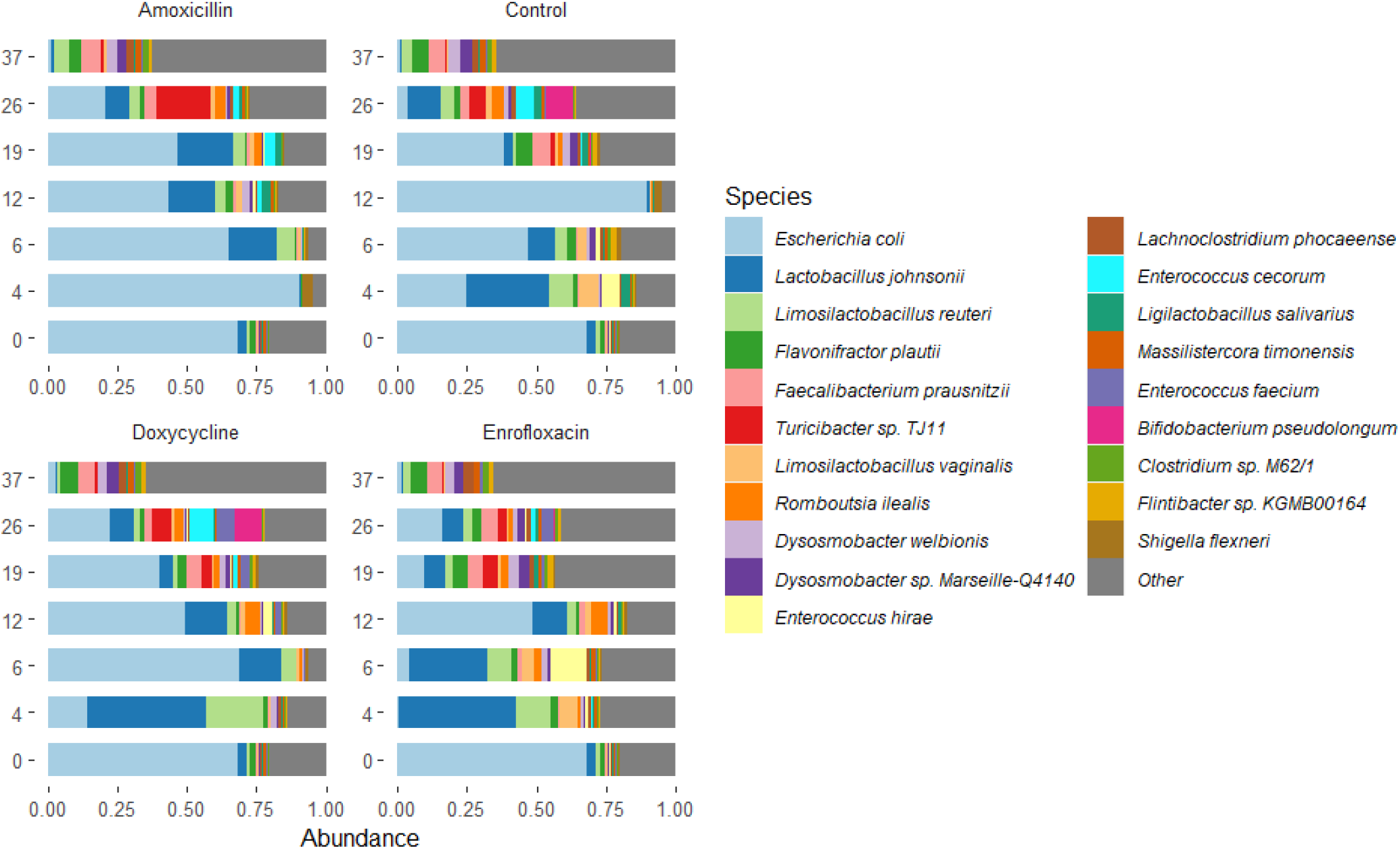
The relative abundance of the samples taken at the different time points after treatment. The 20 most abundance taxa are shown in the legend.

In Figure 5 we determined the alpha diversity of the treated groups at the different time points. For the amoxicillin treatment we found a lower diversity compared to the control group directly after terminating the treatment (p = 0.046). The group treated with enrofloxacin showed a significant difference in the Shannon diversity with the control only at 12 days after start of the treatment (p = 0.045). In contrast, the group treated with doxycycline had an equal alpha diversity to the control. We also investigated beta diversity and we calculated the distance from the baseline sample to the samples taken at other time points as is shown in Figure 6. The group treated with amoxicillin differed from the control group at 4 days after treatment (p = 0.015). Similarly the treatment group of enrofloxacin shows a difference 6 days (p = 0.0062) from beginning the application to the control group. For the group treated with doxycycline we did not find a difference with the control group.

**Figure 5.**
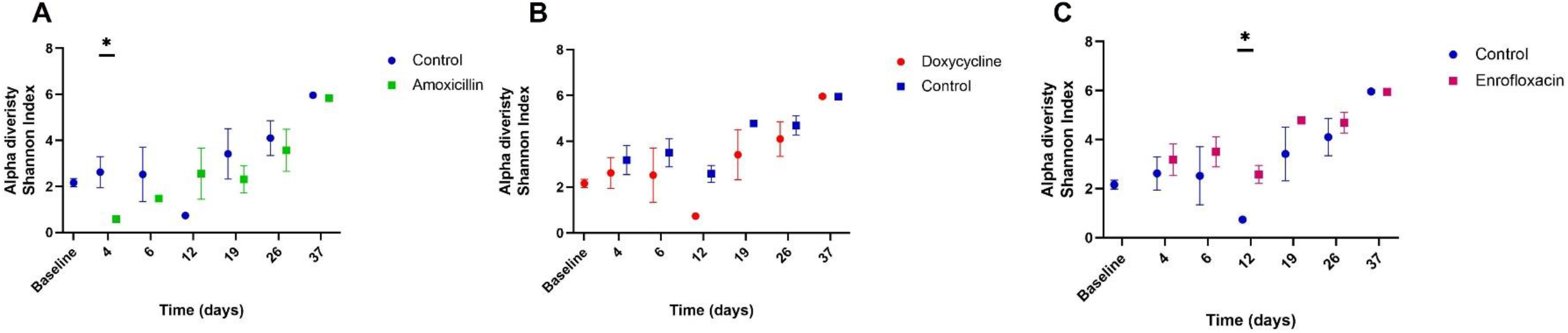
Alpha diversity of the samples at the different time points after start of the application. The alpha diversity is measured in the Shannon index. **A** is the amoxicillin treatment, **B** the doxycycline treatment and **C** the treatment with enrofloxacin. The data points are representing the mean and the standard error. *P≤ 0.05.

**Figure 6.**
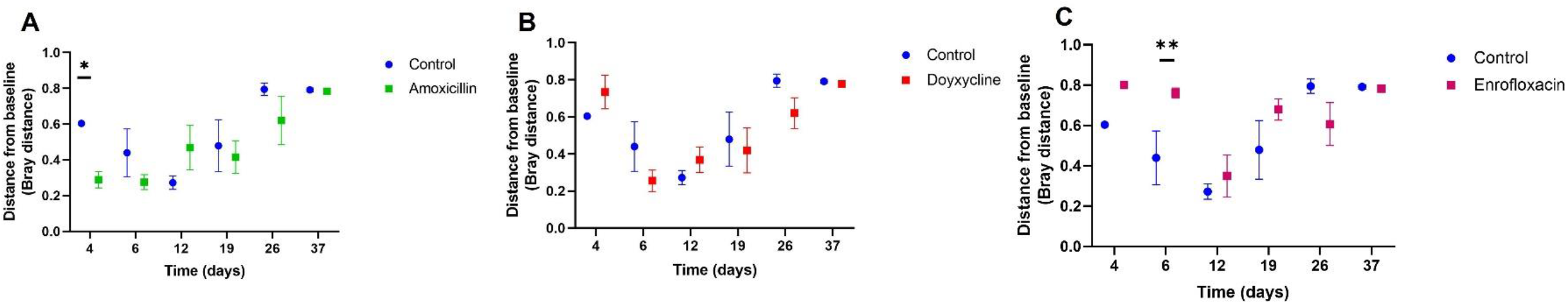
Beta diversity distance of the samples at the different time points after start of the treatment, higher distance equals a more different microbiome. The beta diversity is measured with Curtis-Bray distance and is displayed as distance from the baseline. **A** is the amoxicillin treatment, **B** the doxycycline treatment and **C** the treatment with enrofloxacin. The data points are representing the mean and the standard error. * P ≤ 0.05, ** p ≤ 0.001.

## Discussion

Our findings show that persistent antimicrobial compounds like doxycycline and enrofloxacin remain in the faeces of the broilers for the length of a broiler production round (41 days) at or above MSC level, resulting in prolonged selection pressure for AMR bacteria. Moreover these finding are confirming our hypothesis that both doxycycline and enrofloxacin led to higher levels of resistance than the control group, while amoxicillin did not, even though the effects on the microbiome following amoxicillin treatment were the most pronounced. These results reveal the importance of considering antimicrobial persistence when antimicrobials are priority-classified.

To elaborate, worldwide antimicrobial stewardship programmes are implemented to create awareness for judiciously selecting antimicrobial compounds, for example in livestock. The results of this research showed that persistent antimicrobials, such as doxycycline and enrofloxacin, can retain selective long after cessation of the treatment. Doxycycline from 1.5 to 6 mg/L and enrofloxacin from 1.8 to 15 mg/L within the MSC range. This is in contrast to the non-persistent antimicrobial amoxicillin where hardly no residues were encountered, which is not surprising due to its instability. ^37,38^ One can argue that amoxicillin could be used instead of persistent antimicrobials, however amoxicillin is still used extensively and has selective properties for beta-lactamase genes with possible co-selection for tetracycline resistance. ^39,40^ Nevertheless, persistency of antimicrobials should be taken into consideration in stewardship programmes as it extends the selection phenotypically resistant *E. coli* (doxycycline approximately 52% increase, enrofloxacin approximately 100% increase) and increase the occurrence of resistant genes (doxycycline; significance p ≤ 0.05 and p ≤ 0.0001). Especially in livestock where coprophagic behaviour is not uncommon. For the global efforts to encourage prudent usage of antimicrobials for reducing AMR, persistency would be valuable to include in the classifications of antimicrobials.

Investigating the resistance levels phenotypically and genetically in relation to the persistent residues gives a better understanding about prolonged selection for AMR bacteria due to recirculation of residues. Extensive research has been conducted to resistance levels in the farm environment ^41–43^ but to our knowledge not in combination with antimicrobial residual concentrations. Peng et al. (2016) measured the antimicrobial concentrations of amoxicillin, doxycycline and ciprofloxacin (a fluoroquinolone closely related to enrofloxacin) in manure during and after treatment of caged laying hens. ^44^ In that study birds were administered 50 mg/kg (among different concentrations) for all three the antimicrobials similar to our study. Peng et al. (2016) did find residual concentrations of amoxicillin after treatment, however up to 2 days after the application they could not detect any concentrations in the group treated with 50 mg/kg. This is a slight difference compared to our study since we did not measure any concentrations of amoxicillin directly after treatment. Amoxicillin was probably degraded rapidly by the beta-lactamase enzymes produced by the resistant bacteria or just by the general instability of the compound. This difference could be explained by the beta-lactamase genes present in the microbiome. Peng et al. (2016) did not study the resistome which could influence the amoxicillin concentration. Furthermore, in the study of Peng et al. (2016) doxycycline and ciprofloxacin were measured with similar concentrations as we observed in our study (doxycycline 6 mg/kg and enrofloxacin 15 mg/kg). After the application time Peng et al. (2016) did not detect any doxycycline after 7 days which is unlike our findings as we measured residues after 37 days. Furthermore, ciprofloxacin was measured with an average concentration of 5.78 mg/kg after terminating the treatment, comparable to our enrofloxacin measurements and the same trend as other studies determined. ^45,46^

The objective of this study was to investigate if persistent antimicrobial residues lead to increased AMR levels (genotypically or phenotypically). The Monitoring of Antimicrobial Resistance and Antibiotic Usage in Animals in the Netherlands (MARAN) measured a prevalence of 40% ampicillin resistance in *E. coli* in broilers. ^47^ The presence of amoxicillin resistance in the baseline measurement suggested existing resistance in the broilers, implying that there were already beta-lactam producing bacteria present, despite our efforts to select offspring from a parental flock which was not treated. This may have influenced the amoxicillin treatment group since we observed high amoxicillin resistance phenotypically and genetically directly after the treatment, possibly because of direct selection of resistant bacteria. We encountered higher levels of resistance in the resistome data at day 37 for doxycycline and enrofloxacin. This can be explained as we used caecal material instead of faecal droppings. Caecal samples have been observed to have higher level of resistance genes.^48^ This is explaining the increased resistant levels compared to the control group despite the concentrations which are below the MSC at day 37.

Considering the resistant genes in the resistome sequenced from the enrofloxacin treatment, it can be argued that enrofloxacin does not select for resistant genes. Actually, this is described in the literature; resistance towards enrofloxacin involves another selection mechanism based on selection or induction of single nucleotide polymorphism (SNPs) associated with resistance instead of an increase in species carrying resistance genes. ^49,50^ We sequenced several of the enrofloxacin resistant *E. coli* strains from the enrofloxacin treatment where we observed SNPs responsible for enrofloxacin resistance (unpublished data).

Our experimental set up is different from conditions of commercial broilers, such as temperature or humidity. These parameters would be different and less constant which might influence the results generated from this research. However, we considered the density of the broilers as an essential parameter since this is important for mimicking the commercial setting and also to enable coprophagy. To conclude, we believe that persistency is an important factor in development of AMR.

## Acknowledgements

We would like to thank the animal caretakers from the animal facility Utrecht University (Utrecht, The Netherlands) for their assistance during the animal trial. Besides this we would like to thank Robbert van den Beld of WFSR (Wageningen Food Safety Research, The Netherlands) for his guidance and assistance for extracting the antimicrobials. Lastly we would like to thank André Steentjes of Veterinair Centrum Someren for assisting to select parental flocks of broilers for our trial.

## Transparency declarations

None to declare.

## Supplementary Data

Supplementary materials for this article may be found online at 10.5281/zenodo.10720991.

Sequence data for this article can be found in the SRA under accession PRJEB73721.

